# Occurrence and characterization of cyst nematode species (*Globodera* spp.) associated with potato crops in Colombia

**DOI:** 10.1101/2020.10.13.337238

**Authors:** Daniela Vallejo, Diego A. Rojas, John A. Martinez, Sergio Marchant, Claudia M. Holguin, Olga Y. Pérez

## Abstract

Potato cyst nematodes (PCN) from the genus *Globodera* spp. cause major losses in potato (*Solanum tuberosum*) industry worldwide. Despite their importance, at present little is known about the status of this plant pathogen in cultivated potatoes in Colombia. In this study, a total of 589 samples collected from 75 geographic localities from nine potato producing departments of Colombia were assayed for the presence of potato cyst nematodes. Fifty-seven percent of samples tested positive for PCN. All populations but one were identified as *Globodera pallida*, with conspicuous morphometric variation found among populations. Based on phylogenetic analysis of the internal transcribed spacer region (ITS1-5.8S-ITS2) of the rRNA gene and D2-D3 expansion segments of the 28S rRNA gene, *G. pallida* from Colombia formed a monophyletic group closely related to Peruvian populations, with the lowest average number of nucleotide substitutions per site (*Dxy*= 0.002) and net nucleotide substitutions per site (*Da*= 0.001), when compared to *G. pallida* populations from South, North America and Europe. A single sample formed a well-supported subclade along with *G. rostochiensis* and *G. tabacum* from Japan, USA and Argentina. To our knowledge this is the first comprehensive survey of *Globodera* populations from Colombia that includes morphological and genetic data. Our findings on species diversity and phylogenetic relationships of *Globodera* populations from Colombia may help elucidate the status and distribution of *Globodera* species, and lead to the development of accurate management strategies for the potato cyst nematodes.

## Introduction

The cyst nematodes, *Globodera* Skarbilovich, 1959, are one of the most limiting plant parasitic nematodes around the world [1]. Within the genus, thirteen species have been identified, of which *G. rostochiensis, G. pallida, G. ellingtonae*, and *G. tabacum* are important for agriculture [2]. The potato cyst nematodes (PCN), *Globodera rostochiensis* (golden or yellow potato nematode) and *Globodera pallida* (pale potato nematode) cause major losses in potato (*Solanum tuberosum* L.) crops [3], and are also considered as official control pests in many countries [4]. These species cause damage to the potato plants, by penetrating and feeding into the root tissue, which causes nutritional and water deficiency that is expressed in chlorosis and wilting of the leaves, and may also cause low growth, dwarfism and proliferation of small lateral roots that lead to yield reduction [4]. If PCN species are left uncontrolled may reduce potato yield up to 80% [5,6], representing major economic losses in the potato industry worldwide.

Identification of *Globodera* species based on morphological characterization of the perineal area of cysts (e.g. distance from vulva and anus and Granek’s ratio) and some characters of the second stage juvenile (e.g. stylet length and stylet knob shape) [4,7] may be ambiguous. Morphometric measurements of these characters often show overlap among species, making morphological identification of cyst nematodes time consuming and difficult, especially when differentiating *G. pallida* from *G. rostochiensis* and *G. tabacum* species complex (*G*. *tabacum tabacum, G. tabacum solanacearum* and *G*. *tabacum virginiae*) [8,9]. Therefore, molecular diagnosis is a necessary and recommended complement to identify cyst nematode species [4].

For plant-parasitic nematodes, molecular diagnostics not only improve speed and accuracy of nematode identification, but also have allowed a better understanding of the biology of nematodes as agricultural pests [10]. The genomic regions more often used to study phylogenetic relationships for plant-parasitic nematodes include DNA fragments from the 28S ribosomal DNA (rDNA), internal transcribed spacer (ITS), as well as mitochondrial DNA (mtDNA) [2,10–15]. Ribosomal genes exhibit enough conserved inter-specific neutral genetic variation as to inform species delimitation without being prone to marker saturation [15–18]. For cyst identification, although several methods have been used, DNA-based approaches have shown to be more accurate to separate *G. pallida* from *G. rostochiensis* and other *Globodera* species and, ribosomal regions have also shown to be useful markers to distinguish species within the genus [12,17,19–21]. For new occurrences of *Globodera* spp., sequencing of DNA fragments is also recommended, especially for regions where genetic data has not been reported before and for PCN species that may not follow a typical profile [17,22]. For *Globodera* species from Colombia, genetic information including validation of currently available diagnostic DNA markers and molecular phylogenetics have not been documented.

In Colombia, *G. pallida* was first identified based on morphological characters in 1970 in Cumbal, municipality located in the Nariño department, at the south west extreme of the country [23]. In 1971, the species was regulated under the authority of The Instituto Colombiano Agropecuario (ICA) and listed as quarantine pest, limiting the access to export potato seeds from Nariño and its neighbor department, Cauca, to other producing potato departments of Colombia. In 1983, Nieto [24] conducted an intensive PCN survey and reported *G. pallida* in other municipalities of Nariño (Túquerres, Pupiales, Ipiales, Gualmatán, Sapuyes, among others), as well as in Cauca (Totoró, Cajibío, Silvia, Popayán, Páez, among others), with an average of 50-80 cysts/100 g of soil in Nariño and 9-10 cysts/100 g of soil in Cauca. The authors also sampled the nematode in Cundinamarca and Boyacá, the main producing potato departments in Colombia, and other minor producing potato departments such as Caldas, Tolima, Valle del Cauca, Santander and Norte de Santander, but only reported the presence of PCN in Nariño and Cauca. In 2004, the species was no longer listed as an official control pest. Yet, in a survey conducted from 2011 to 2012, PCN was reported in 12 out of 14 sampled fields in Tunja, Samacá and Ventaquemada (municipalities of Boyacá) and Tausa, Tabio and Zipaquirá (municipalities of Cundinamarca), although population densities were not registered [25]. Therefore, PCN was considered as re-emerging pathogen in 2012 by the Federación Colombiana de Productores de Papa (Colombian Federation of Potato Producers – FEDEPAPA), and ICA [25].

To obtain better knowledge about *Globodera* spp. associated with potato crops in Colombia, it is necessary to develop DNA sequence information to better characterize populations from different geographic regions and to understand their distribution patterns. This information will also serve as a foundation to the design of effective control measures that require fast and accurate identification of species, and it is a crucial factor when searching for possible sources of host-plant resistance as well as for other management strategies. Therefore, the objectives of this study were to: i) survey the *Globodera* spp. populations detected in cultivated potatoes in Colombia; ii) carry out a molecular characterization of these *Globodera* populations based on sequences of the ITS1 of rRNA, partial 18S rRNA and, D2-D3 expansion segments of the 28S nuclear ribosomal RNA gene; iii) study the phylogenetic relationships of *Globodera* spp. from Colombia by comparison with previously published molecular data of populations from other regions of the world; and iv) compare cyst morphometric measurements among *Globodera* populations from Colombia and other species previously reported.

## Materials and Methods

### Ethics statement

Nematode sampling was performed under a collection permit granted by the Autoridad Nacional de Licencias Ambientales (ANLA) [Colombian National Authority Environmental Permits]: “Permit for collecting specimens of wild species of the biological diversity for non-commercial scientific research purposes], resolution No. 1466, expedited on December 3, 2014.

### Nematode populations and sampling

From 2013 to 2017, an extensive survey was conducted throughout the main commercial potato producing regions of Colombia. A total of 589 sampling sites were selected in 75 municipalities using a stratified sampling strategy. The strata were defined as the departments with the highest potato area reported in Colombia [26], for a total of nine departments sampled: Cundinamarca, Boyacá, Antioquia, Nariño, Santander, Norte de Santander, Tolima, Caldas and Cauca (Fig 1, Table 1). At each department, the number of fields sampled per municipality was proportional to the potato area planted and fields at each municipality were selected based on established potato crops in pre-flowering and flowering stages (Fig 1, Table 1). Soil samples at each field were collected from within rows, at roughly equal intervals in a line transect pattern across an area of 10,000 m^2^ or less. A soil sample consisted of 60 soil cores (1.5 cm in diameter by 5 cm deep) taken near the root of the plants. Infected roots and surrounding soil of samples collected from each field were pooled into one composite sample. Samples were placed into plastic bags, transported to the laboratory of microbiology at the Corporación Colombiana de Investigación Agropecuaria (AGROSAVIA), Tibaitatá Research Center, in Mosquera, Cundinamarca, and stored at 4°C until processing. Cyst nematodes were extracted from soil samples using the Fenwick method [27], and cyst individuals per 100 cm^3^ of soil were counted and morphologically identified using the keys by Golden and Handoo et al. [7,28]. Additionally, a viability test was performed by randomly selecting 10 cyst per population that were crushed using a huijsman homogenizer [29] to release eggs and juveniles, alive eggs and j1 were counted under the stereoscope and viability percentage was calculated per population.

**Fig 1.**
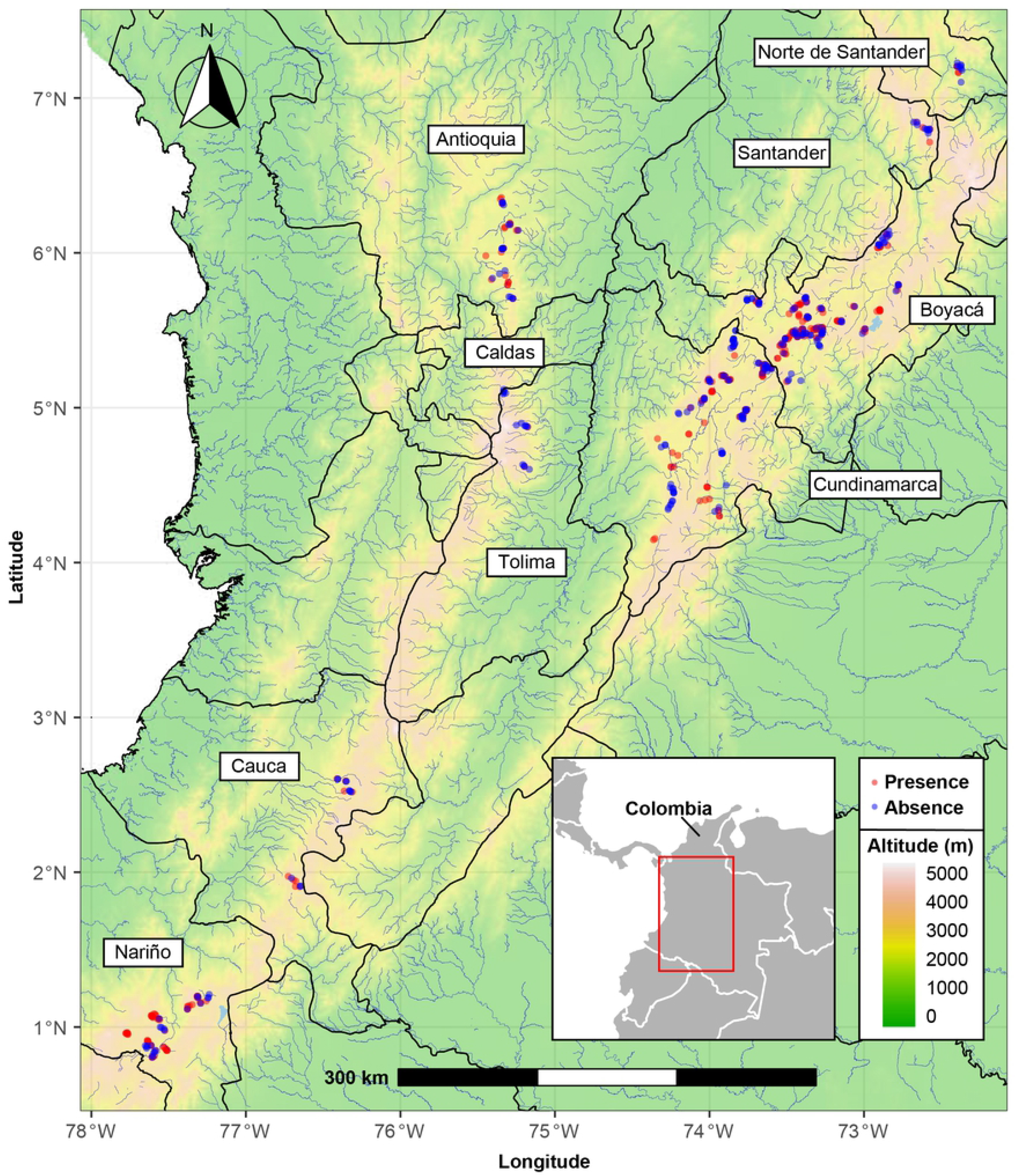
Map of the Colombian Andes showing sampling sites of PCN associated with potato crops in Colombia. Black lines represent department limits, blue lines represent rivers and lakes. Colors according elevation map. Red dots mark the position of the sample sites that tested positive for PCN and blue dots mark the position of the sampling sites that tested negative for PCN.

**Table 1.** *Globodera* species and density levels (number of individuals per 100 cm^3^ of soil) and prevalence (%) in cultivated potato crops in Colombia.

| Department | Municipality | Species | # of samples collected | # of samples with *Globodera* spp. | Prevalence | Average/100 cm<sup>3</sup> soil | Population range/100 cm<sup>3</sup> soil | Viability average | Potato Variety |
| --- | --- | --- | --- | --- | --- | --- | --- | --- | --- |
|  |  |  |  |  | (%) |  |  | (%) |  |
| Nariño | Túquerres | *G. pallida* | 12 | 12 | 100 | 97,3 | 1 - 345 | 60 | D_Capiro |
| Nariño | Ipiales | *G. pallida* | 10 | 5 | 50 | 9,1 | 2 - 45 | 41 | D_Capiro |
| Nariño | Pasto | *G. pallida* | 15 | 9 | 60 | 14,7 | 1 - 64 | 60 | D_Capiro |
| Nariño | Yacuanquer | *G. pallida* | 2 | 2 | 100 | 3,5 | 3 - 4 | 45 | Criolla |
| Nariño | Tangua | *G. pallida* | 5 | 4 | 80 | 11 | 1 - 34 | 55 | D_Capiro |
| Nariño | Ospina | *G. pallida* | 5 | 4 | 80 | 47,4 | 5 - 93 | 38 | D_Capiro |
| Nariño | Iles | *G. pallida* | 5 | 1 | 20 | 16 | 1 - 80 | 59 | D_Capiro |
| Nariño | Guachucal | *G. pallida* | 7 | 7 | 100 | 21,3 | 1 - 84 | 63 | Betina |
| Nariño | Pupiales | *G. pallida* | 11 | 6 | 54,5 | 5,9 | 1 - 29 | 30 | D_Capiro |
| Nariño | Córdoba | *G. pallida* | 7 | 7 | 100 | 9,4 | 1 - 27 | 44 | P Suprema |
| Cundinamarca | Chipaqué | *G. pallida* | 4 | 4 | 100 | 68,3 | 2-168 | 51 | P Suprema |
| Cundinamarca | Madrid | *G. pallida* | 5 | 2 | 40 | 1,8 | 2 - 7 | 63 | D_Capiro |
| Cundinamarca | Pasca | - | 6 | 0 | 0 | 0 | 0 | 0 | P Suprema |
| Cundinamarca | Sesquilé | *G. pallida* | 9 | 5 | 55,6 | 10,2 | 1 - 25 | 62 | P Suprema |
| Cundinamarca | Subachoque | *G. pallida* | 5 | 1 | 20 | 4 | 0 - 20 | 52 | P Suprema |
| Cundinamarca | Tausa | *G. pallida* | 25 | 17 | 68 | 84,2 | 1 - 1327 | 51 | P Suprema |
| Cundinamarca | Ubaque | *G. pallida* | 4 | 2 | 50 | 1,8 | 3 - 4 | 58 | P Superior |
| Cundinamarca | Ubaté | *G. pallida* | 3 | 3 | 100 | 443,3 | 268 - 556 | 51 | P_Pastusa |
| Cundinamarca | Une | *G. pallida* | 3 | 3 | 100 | 3,3 | 1 - 5 | 72 | Criolla |
| Cundinamarca | Villapinzón | *G. pallida* | 29 | 18 | 62,1 | 23,4 | 1 - 475 | 74 | P Suprema |
| Cundinamarca | Zipacquirá | *G. pallida* | 9 | 5 | 55,6 | 2 | 2 - 10 | 64 | P Suprema |
| Cundinamarca | Cajicá | *G. pallida* | 1 | 1 | 100 | 1 | 0 - 1 | 80 | D_Capiro |
| Cundinamarca | Tenjo | *G. pallida* | 2 | 2 | 100 | 72,5 | 50-95 | 76 | D_Capiro |
| Cundinamarca | Sibaté | - | 8 | 0 | 0 | 0 | 0 | 0 | Criolla |
| Cundinamarca | Soacha | *G. pallida* | 3 | 3 | 100 | 1 | 0 - 1 | 76 | D_Capiro |
| Cundinamarca | Cogua | *G. pallida* | 4 | 4 | 100 | 1,5 | 1 - 2 | 73 | Betina |
| Cundinamarca | San Bernardo | *G. pallida* | 2 | 2 | 100 | 2,5 | 2 - 3 | 51 | Criolla |
| Cundinamarca | Fosca | *G. pallida* | 4 | 2 | 50 | 2 | 0 - 5 | 69 | P Suprema |
| Cundinamarca | Lenguazaque | *G. pallida* | 3 | 1 | 33,3 | 1,7 | 0 | 60 | P_Pastusa |
| Cundinamarca | Simijaca | - | 2 | 0 | 0 | 0 | 0 - 40 | 0 | P Suprema |
| Cundinamarca | Susa | *G. pallida* | 15 | 1 | 6,7 | 2,7 | 1 - 125 | 55 | P Suprema |
| Cundinamarca | Guatavita | *G. pallida* | 11 | 4 | 36,4 | 11,8 | 1 - 125 | 55 | P Suprema |
| Cundinamarca | La Calera | - | 5 | 0 | 0 | 0 | 0 | 0 | P Suprema |
| Cundinamarca | Choconta | *G. sp.* | 8 | 7 | 87,5 | 43 | 1 - 93 | 70 | P Suprema |
| Boyacá | Tota | *G. pallida* | 6 | 4 | 66,7 | 4,7 | 2 - 19 | 58 | P_Pastusa |
| Boyacá | Toca | *G. pallida* | 10 | 7 | 70 | 27,3 | 5 - 122 | 51 | Tocarreña |
| Boyaca | Tunja | *G. pallida* | 25 | 17 | 68 | 47,3 | 1 - 411 | 60 | D_Capiro |
| Boyaca | Chíquiza | *G. pallida* | 11 | 9 | 81,8 | 47 | 1 - 142 | 23 | Betina |
| Boyaca | Úmbita | *G. pallida* | 5 | 1 | 20 | 2 | 0 - 2 | 20 | Tocarreña |
| Boyaca | Samacá | *G. pallida* | 29 | 22 | 75,9 | 53,2 | 1 - 429 | 24 | D_Capiro |
| Boyaca | Oicatá | *G. pallida* | 6 | 4 | 66,7 | 7,7 | 1 - 36 | 24 | D_Capiro |
| Boyaca | Sogamoso | *G. pallida* | 8 | 8 | 100 | 107,5 | 1 - 305 | 38 | P_Pastusa |
| Boyaca | Siachoque | *G. pallida* | 27 | 20 | 74,1 | 64 | 1 - 364 | 54 | P_Pastusa |
| Boyaca | Arcabuco | *G. pallida* | 10 | 5 | 50 | 122,5 | 1 - 873 | 23 | P_Pastusa |
| Boyaca | Ventaquemada | *G. pallida* | 15 | 12 | 80 | 7,9 | 1 - 73 | 53 | P Suprema |
| Boyaca | Motavita | *G. pallida* | 8 | 4 | 50 | 2,6 | 1 - 14 | 54 | Betina |
| Boyaca | Sora | *G. pallida* | 3 | 3 | 100 | 185 | 101 - 286 | 60 | D_Capiro |
| Boyaca | Saboyá | *G. pallida* | 16 | 5 | 31,3 | 1,4 | 1 - 15 | 36 | P_Pastusa |
| Boyaca | Soracá | *G. pallida* | 9 | 8 | 88,9 | 84,7 | 1 - 306 | 24 | D_Capiro |
| Boyaca | Boyacá | *G. pallida* | 3 | 1 | 33,3 | 13,7 | 1 - 41 | 35 | D_Capiro |
| Boyaca | Viracachá | *G. pallida* | 3 | 2 | 66,7 | 1,3 | 1 - 2 | 32 | Rubí |
| Boyaca | Ciénega | - | 4 | 0 | 0 | 0 | 0 | 0 | P Suprema |
| Boyaca | Mongua | *G. pallida* | 2 | 1 | 50 | 0,5 | 0 - 1 | 38 | ICA_Única |
| Boyaca | Firavitoba | *G. pallida* | 3 | 2 | 66,7 | 1 | 1 - 2 | 44 | P_Pastusa |
| Boyaca | Gámeza | *G. pallida* | 5 | 2 | 40 | 0,8 | 1 - 2 | 20 | Tocarreña |
| Boyaca | Belén | *G. pallida* | 11 | 7 | 63,6 | 122,3 | 1 - 757 | 40 | P_Pastusa |
| Boyaca | Tutazá | *G. pallida* | 12 | 3 | 25 | 6,9 | 1 - 78 | 49 | ICA_Única |
| Cauca | San Sebastián | *G. pallida* | 8 | 5 | 62,5 | 4,1 | 1 - 13 | 48 | P_Pastusa |
| Cauca | Silvia | *G. pallida* | 6 | 2 | 33,3 | 0,3 | 0 - 1 | 47 | Criolla |
| Cauca | Totoró | *G. pallida* | 6 | 2 | 33,3 | 4,3 | 1 - 23 | 51 | Criolla |
| Antioquia | San Vicente | *G. pallida* | 9 | 5 | 55,6 | 7,11 | 2 - 24 | 40 | Criolla |
| Antioquia | La Unión | *G. pallida* | 9 | 3 | 33,3 | 6 | 1 - 51 | 32 | D_Capiro |
| Antioquia | Sonsón | *G. pallida* | 9 | 5 | 55,6 | 7,5 | 1 - 29 | 16 | Criolla |
| Antioquia | La Ceja | *G. pallida* | 1 | 1 | 100 | 7 |  | 20 |  |
| Antioquia | Abejorral | *G. pallida* | 6 | 3 | 50 | 2,17 | 1 - 7 | 15 | Criolla |
| Antioquia | Marinilla | *G. pallida* | 7 | 5 | 71,4 | 5,71 | 1 -26 | 20 | Criolla |
| Antioquia | Santuario | *G. pallida* | 3 | 2 | 66,7 | 1,33 | 1 - 3 | 20 | ICA_Nevada |
| Norte de Santander | Pamplona | *G. pallida* | 8 | 2 | 25 | 0,9 | 1 - 4 | 37 | Criolla |
| Norte de Santander | Cácota | *G. pallida* | 4 | 2 | 50 | 1 | 1 - 2 | 16 | Criolla |
| Santander | Concepción | *G. pallida* | 8 | 2 | 25 | 0,9 | 1 - 6 | 20 | P_Pastusa |
| Santander | Carcasí | *G. pallida* | 2 | 1 | 50 | 0,5 | 0 - 1 | 36 | P_Pastusa |
| Santander | Cerrito | *G. pallida* | 4 | 1 | 25 | 1 | 1 - 4 | 31 | ICA_Única |
| Caldas | Manizales | *G. pallida* | 7 | 1 | 14,3 | 0,43 | 1 - 3 | 15 | P_Pastusa |
| Tolima | Anzoátegui | *G. pallida* | 5 | 1 | 20 | 0,4 | 1 - 2 | 29 | P Suprema |
| Tolima | Murillo | *G. pallida* | 7 | 1 | 14,3 | 0,2 | 0 - 1 | 19 | P_Pastusa |

### DNA extraction, Polymerase Chain Reaction and sequencing

For molecular characterization, from the departments that showed the highest cyst nematode densities, soil samples taken per municipality were pooled according to proximity distance (1 km and 5 km range), resulting in one to two samples per municipality, for a total of 26 populations analyzed (Table 1). From each pooled population, DNA was extracted from individual cysts or juveniles using the “Sigma Extract-N-Amp Kit (XNAT2)” kit (Sigma, St. Louis, MO) according to the protocol reported by Ma et al. (2011) at AGROSAVIA, La Selva Research Center, Rionegro, Antioquia. The number of individuals analyzed per population depended upon the cyst nematode density present in each soil sample. DNA was then stored at - 20°C until used.

PCR amplification of two genomic regions were performed using 12.5 μl of the Extract-N-AmpTM Tissue PCR kit (Sigma), 1 μl of each primer, 4 μl of DNA and water to complete a volume of 25 μl. The rDNA primers used for PCR and DNA sequencing are listed in Table 2. The ITS region of ribosomal DNA was amplified using 94° C for 2.5 min for initial denaturation, followed by 40 cycles at 94° C for 1 min, 55° C for 1 min, 72° C for 2 min, and a final extension of 72° C for 5 minutes. For the 28S region, initial denaturation was 94° C for 5 min, 40 cycles of 94° C for 30 sec, 58° C for 30 sec, 72° C for 1 min, and a final extension of 72° C for 10 min (Nunn, 1992). The products were loaded on a 1.5% agarose gel and visualized using gel red (Biotium, San Francisco, CA). Sanger sequencing of the amplicons was performed in both directions by CorpoGen (Bogotá, Colombia).

**Table 2.**
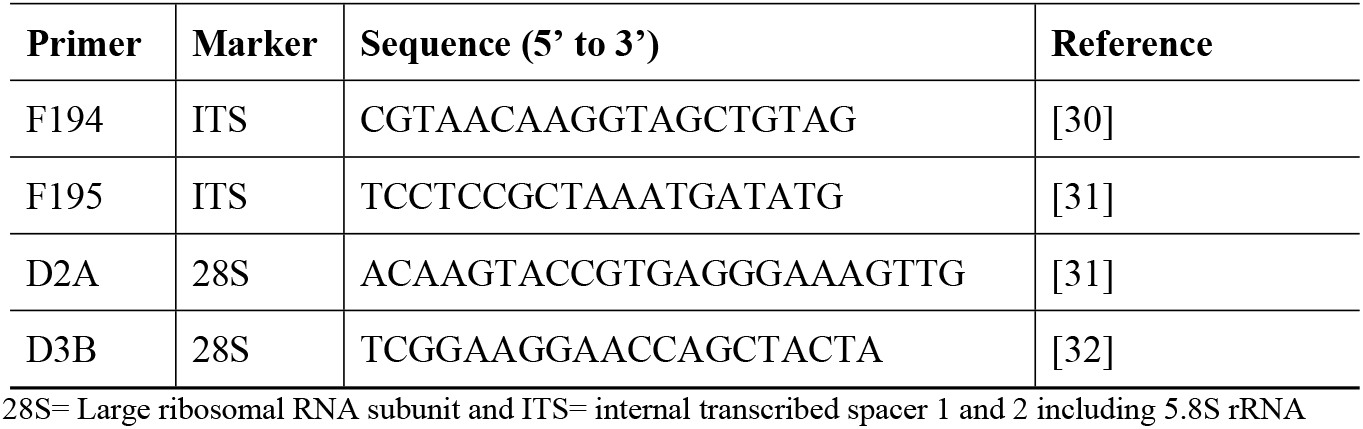
Primers used for polymerase chain reaction and DNA sequencing of *Globodera* spp. individuals recovered from cultivated potatoes in Colombia.

### Sequence alignment and Phylogenetic analyses

Resulting sequences were assembled in Sequencher^®^ software version 5.1 (Gene Codes Corporation, Ann Arbor, MI USA) and manually reviewed for base calling errors. Partial 28S rRNA and ITS1-2 + 5.8S rRNA gene sequences from *G. pallida, G. mexicana, G. rostochiensis, G. tabacum, G. ellingtonae*, and *G. artemisiae*, were retrieved from GenBank nucleotide database and included in the alignment (Table 3) [8,19,21,33–42]. Sequences of *Punctodera punctata* and *P. chalcoensis* also obtained from GenBank (AF274416.1, DQ328699.1.1, AY090885. 1), were used as outgroup taxa for both gene regions. After that, sequence alignments were performed using Clustal W [43] and manually edited using Gene Doc v.2.7 [44]. To remove ambiguous regions in the alignment the program Gblocks was used with the standard settings [45]. Newly generated sequences for both gene regions were deposited in GenBank (Table 3).

**Table 3.** Details of cyst nematode populations included in the molecular and phylogenetic studies from cultivated potatoes in Colombia and reported in other studies.

| *Globodera* species | Location<br>(Municipality, Department) | Specimen code | Accession number 28S | Accession number ITS |
| --- | --- | --- | --- | --- |
| *Globodera pallida* | La Unión, AN* | A1A | MH389939 | MH389979 |
| *Globodera pallida* | La Unión, AN | A1B | MH389938 | MH389978 |
| *Globodera pallida* | La Unión, AN | A1C | - | MH389977 |
| *Globodera pallida* | Ventaquemada, BO | B2A | MH389946 | MH389986 |
| *Globodera pallida* | Ventaquemada, BO | B2B | MH389945 | MH389985 |
| *Globodera pallida* | Ventaquemada, BO | B2C | MH389944 | MH389984 |
| *Globodera pallida* | Ventaquemada, BO | B2D | MH389943 | MH389983 |
| *Globodera pallida* | Arcabuco, BO | B3D | MH389942 | MH389982 |
| *Globodera pallida* | Chíquiza, BO | B4D | MH389937 | - |
| *Globodera pallida* | Tunja, BO | B5D | - | MH389976 |
| *Globodera pallida* | Toca, BO | B6D | MH389936 | MH389975 |
| *Globodera pallida* | Sogamoso, BO | B7A | MH389935 | MH389974 |
| *Globodera pallida* | Sogamoso, BO | B7B | MH389934 | - |
| *Globodera pallida* | Sogamoso, BO | B7D | MH389933 | - |
| *Globodera pallida* | Sora, BO | B8A | MH389932 | MH389973 |
| *Globodera pallida* | Sora, BO | B8C |  | MH389972 |
| *Globodera pallida* | Sora, BO | B8D |  | MH389971 |
| *Globodera pallida* | Samacá, BO | B9D | MH389941 | MH389981 |
| *Globodera pallida* | Soracá, BO | B11A | MH389931 |  |
| *Globodera pallida* | Soracá, BO | B11B | MH389930 |  |
| *Globodera pallida* | Soracá, BO | B11C | MH389929 |  |
| *Globodera pallida* | Soracá, BO | B12A | MH389940 | MH389980 |
| *Globodera pallida* | Susa, CU | C1A | MH389928 | MH389970 |
| *Globodera pallida* | Susa, CU | C1B | MH389927 |  |
| *Globodera pallida* | Susa, CU | C1D |  | MH389969 |
| *Globodera pallida* | Guatavita, CU | C2B | MH389926 | MH389968 |
| *Globodera pallida* | Guatavita, CU | C2C | MH389925 | MH389967 |
| *Globodera pallida* | Guatavita, CU | C3C | MH389923 | MH389965 |
| *Globodera pallida* | Guatavita, CU | C3D | MH389922 |  |
| *Globodera pallida* | Ubaté, CU | C4C | MH389921 |  |
| *Globodera pallida* | Ubaté, CU | C4D | MH389920 |  |
| *Globodera pallida* | Tausa, CU | C5A | MH389919 | MH389964 |
| *Globodera pallida* | Tausa, CU | C5B | MH389918 |  |
| *Globodera pallida* | Subachoque, CU | C6B | MH389917 |  |
| *Globodera pallida* | Sesquilé, CU | C8A | MH389916 |  |
| *Globodera pallida* | Sesquilé, CU | C8C | MH389915 | MH389963 |
| *Globodera pallida* | Sesquilé, CU | C8D |  | MH389962 |
| *Globodera pallida* | Cajicá, CU | C10D |  | MH389961 |
| *Globodera pallida* | Villapinzón, CU | C11A | MH389914 | MH389960 |
| *Globodera pallida* | Villapinzón, CU | C11D | MH389913 | MH389959 |
| *Globodera pallida* | Túquerres, NA | N2A | MH389949 | MH389990 |
| *Globodera pallida* | Túquerres, NA | N2B | MH389948 | MH389989 |
| *Globodera pallida* | Túquerres, NA | N2C |  | MH389988 |
| *Globodera pallida* | Túquerres, NA | N2D | MH389947 | MH389987 |
| *Globodera pallida* | Guachucal, NA | N4A | MH389912 |  |
| *Globodera pallida* | Guachucal, NA | N4B | MH389911 |  |
| *Globodera pallida* | Guachucal, NA | N4C | MH389910 | MH389958 |
| *Globodera pallida* | Guachucal, NA | N4D | MH389909 |  |
| *Globodera pallida* | Belén, NA | N5A |  | MH389957 |
| *Globodera pallida* | Belén, NA | N5B |  | MH389956 |
| *Globodera pallida* | Belén, NA | N5C | MH389908 | MH389955 |
| *Globodera pallida* | Belén, NA | N5D | MH389907 | MH389954 |
| *Globodera pallida* | Ospina, NA | N8B |  | MH389953 |
| *Globodera pallida* | Ospina, NA | N8C | MH389906 | MH389952 |
| *Globodera pallida* | Ospina, NA | N8D | MH389905 | MH389951 |
| *Globodera pallida* | Ipiales, NA | N9A | MH389904 | MH389950 |
| *Globodera sp* | Chocontá, CU | C3A | MH389924 | MH389966 |
| *Globodera pallida* | Peru |  |  | GU084813.1 |
| *Globodera pallida* | Peru |  |  | GU084805.1 |
| *Globodera pallida* | Chile |  |  | GU084800.1 |
| *Globodera pallida* | Ukraine |  |  | AJ606687.1 |
| *Globodera pallida* | England |  |  | DQ847110.1 |
| *Globodera pallida* | Poland |  |  | EU855119.1 |
| *Globodera pallida* | Peru |  |  | GU084804.1 |
| *Globodera pallida* | Peru |  |  | GU084806.1 |
| *Globodera pallida* | Peru |  |  | HQ670269.1 |
| *Globodera mexicana* | Mexico |  |  | EU006707.1 |
| *Globodera mexicana* | Mexico |  |  | EU006708.1 |
| *Globodera rostochiensis* | USA |  |  | EF153839.1 |
| *Globodera rostochiensis* | Australia |  |  | EF622524.1 |
| *Globodera rostochiensis* | Canada |  |  | FJ212166.1 |
| *Globodera rostochiensis* | England |  |  | EF153840.1 |
| *Globodera rostochiensis* | Bolivia |  |  | GU084809.1 |
| *Globodera tabacum* | Japan |  |  | AB207272.1 |
| *Globodera tabacum* | USA |  |  | GQ294525.1 |
| *Globodera tabacum* | USA |  |  | DQ847112.1 |
| *Globodera tabacum* | Argentina |  |  | DQ097515.2 |
| *Globodera ellingtonae* | USA |  |  | GQ896543.1 |
| *Globodera ellingtonae* | Chile |  |  | GU084808.1 |
| *Globodera ellingtonae* | USA |  |  | DQ097514.2 |
| *Punctodera punctata* | Belgium |  |  | AF274416.1 |
| *Punctodera chalcoensis* | Mexico |  |  | AY090885.1 |
| *Globodera pallida* | France |  | KJ409636.1 |  |
| *Globodera pallida* | Slovakia |  | KJ409626.1 |  |
| *Globodera pallida* | England |  | JN712219.1 |  |
| *Globodera pallida* | Chile |  | JN712220.1 |  |
| *Globodera pallida* | France |  | GU338021.1 |  |
| *Globodera ellingtonae* | USA |  | JN712217.1 |  |
| *Globodera tabacum* | USA |  | GQ294492.1 |  |
| *Globodera rostochiensis* | Canada |  | JN712223.1 |  |
| *Globodera rostochiensis* | Slovakia |  | KJ409625.1 |  |
| *Globodera artemisiae* | Hungary |  | KU845472.1 |  |

Phylogenetic relationships among partial sequences of the 28S rRNA and Internal Transcribed Spacer 1 and 2 plus 5.8S rRNA genes were inferred using Bayesian Inference (BI) and Maximum Likelihood (ML) methods. For both gene regions, the best sequence partition strategy was identified with the Bayesian Information Criterion in PartitionFinder v.2.0 [46]. The sequence partition for 28S rRNA gene was a single partition with HKY substitution model [47] for each of the 3 positions of the codon. The best sequence partition for ITS1-2 + 5.8S rRNA gene internal transcribed spacer 1 and 2 including the 5.8S rRNA region was a partition that included the first and second position of the codon with a K80 model [48] substitution model, and a second partition that consisted on the third codon position under the K80 with proportions of invariable sites (K80+I). Bayesian Inference analyses were performed using MrBayes v.3.1.2 [49], with five independent runs of four Markov chains for 1 x 10^6^ generations and default heating values, sampling every 100 generations with 2500 samples discarded as burn-in after checking for convergence. Clades were considered strongly supported when values were > 0.95 [50]. For Maximum likelihood analyses the software GARLI v.2.0 [51] was used. Bootstrap support for trees was generated with 1,000 replicate searches and summarized in a consensus tree using SumTrees [52], clades were considered as well/strongly supported when bootstrap was >70%. In addition, in order to characterize the genetic divergence between cyst nematodes from Colombia and *Globodera* species already reported, the average number of nucleotide substitution per site (*Dxy*), net nucleotide substitutions per site (*Da*) and number of fixed differences (*Fd*) among genetic groups were computed using DnaSP v.6.11.01 [53].

### Morphometric characterization

For morphometric characterization, 8 to ten cyst nematodes were taken from each of the 26 pooled populations obtained for molecular analysis. Cysts were cleaned and fixed in formaldehyde at 3% allowing preservation in glycerin [7,28]. Then, cysts and vulval cones were photographed and measured using image visualization software (iSolution lite; IMT i-Solution). Morphometric measurements included body width (BW), body length excluding neck (BL), L/W ratio, distance from anus to the nearest edge from fenestra (DAF), fenestra length (FL), Granek’s ratio (GR), and number of cuticular ridges between vulva and anus (NCR) [7,28]. One-way Analysis of variance (ANOVA) was performed on each morphometric variable. The Tukey Studentized Range HDS (Honest Significant Difference) test was used to determine significant differences among population means on the different morphometric measurements at *P* ≤ 0.05.

A linear discriminant analysis (LDA) and a stepwise discriminant analysis (SDA) were also used to determine the best combination of variables that would separate populations based on morphological features. These methods derive a linear combination of variables that summarize between-class variation. The variables included in the initial function for both methods were BW, BL, DAF, FL, NCR, LWR, and GR. The pooled within canonical structure and pooled within class standardized canonical coefficients from the SDA were used to determine each variable’s contribution to the discriminant function. The linear and stepwise discriminant analyses were performed with JMP 14.0.0 (SAS Institute Inc., Cary, NC).

## Results

### Field survey

Of the 589 potato fields sampled, cyst nematodes were detected in 355 fields distributed in 69 municipalities of Colombia, with densities ranging from 1 to 1,327 cysts per 100 cm^3^ of soil (Table 1). The predominant species was *G. pallida*, identified in 51% of the fields sampled in Cundinamarca, 63.3% in Boyacá, 72.2% in Nariño, 54% in Antioquia, 45% in Cauca, 33% in Norte de Santander, 29% in Santander, 17% in Tolima and 14% in Caldas, with densities ranging from 1 to 1327 cysts/100 cm^3^ of soil, 1 to 873 cysts/100 cm^3^ of soil, 1 to 345 cysts/100 cm^3^ of soil, 1 to 51 cysts/100 cm^3^ of soil, 1 to 23 cysts/100 cm^3^ of soil, 1 to 4 cysts/100 cm^3^ of soil, 1 to 6 cysts/100 cm^3^ of soil, 1 to 3 cysts/100 cm^3^ of soil and 1 to 2 cysts/100 cm^3^ of soil, respectively. Among municipalities, the highest mean densities were detected in Ubaté (443.3 cysts/100 cm^3^ of soil), followed by Sora, Arcabuco, Belén, Sogamoso, Túquerres and Tausa (185, 122.5, 122.3, 107.5, 97.3 and 84,2 cysts/100 cm^3^ of soil, respectively). Cyst nematodes were not found in Pasca, Sibaté, Simijaca, and La Calera in Cundinamarca, nor in Ciénega in Boyacá (Table 1). All samples positive for PCN showed cysts with viable eggs, and viability percentage ranged from 15 to 80%. Boyacá and Cundinamarca were the departments that had in average the higher viability (Table 1).

Given host-based grouping, PCN was detected in samples taken from varieties of *Solanum tuberosum* Group *Andigena* such as Diacol Capiro (107 out of 158 samples), Betina (24 out of 30 samples), Pastusa Suprema (81 out of 167 samples), Tocarreña (10 out of 20 samples), Rubí (2 out of 3 samples), ICA Única (4 out of 16 samples), ICA Nevada (2 out of 3 samples) and, from varieties of *Solanum tuberosum* Group *Phureja* such as Criolla variety (32 out of 69 samples) (Table 1). A morphologically different species (under description), was only detected in one field in Chocontá (Cundinamarca) on Suprema variety.

### Molecular analysis

The amplification of D2-D3 expansion segments of 28S rRNA and internal transcribed spacer 1, 2 including the 5.8S rRNA yielded single fragments of 609 and 848 bp, respectively. Forty-two new D2-D3 of 28S rRNA gene sequences and twenty-eight new internal transcribed spacer 1 and 2 including the 5.8S rRNA were obtained in the present study (Table 2).

Phylogenetic relationships inferred from analyses of D2-D3 expansion segments of 28S rRNA of a multiple-edited alignment (57 sequences), showed two well supported major clades based on BI and ML inferences (PP= 1.00, BP= 90) (Fig 2). A highly supported clade (i) (PP= 1.00, BP= 90), was formed by sequences of *G. pallida* from France, Chile, England, Slovakia and all but one cyst nematode sequence obtained in this study from Colombia. The second major clade, Clade (ii) grouped three species, *G. ellingtonae* and *G. tabacum* that formed a well-supported subclade (PP=1.00, BP=83) clearly separated from a politomy formed by a single cyst nematode sample (*G*. sp) from Colombia and *G. rostochiensis* from Canada and Slovakia (Fig 2).

**Fig 2.**
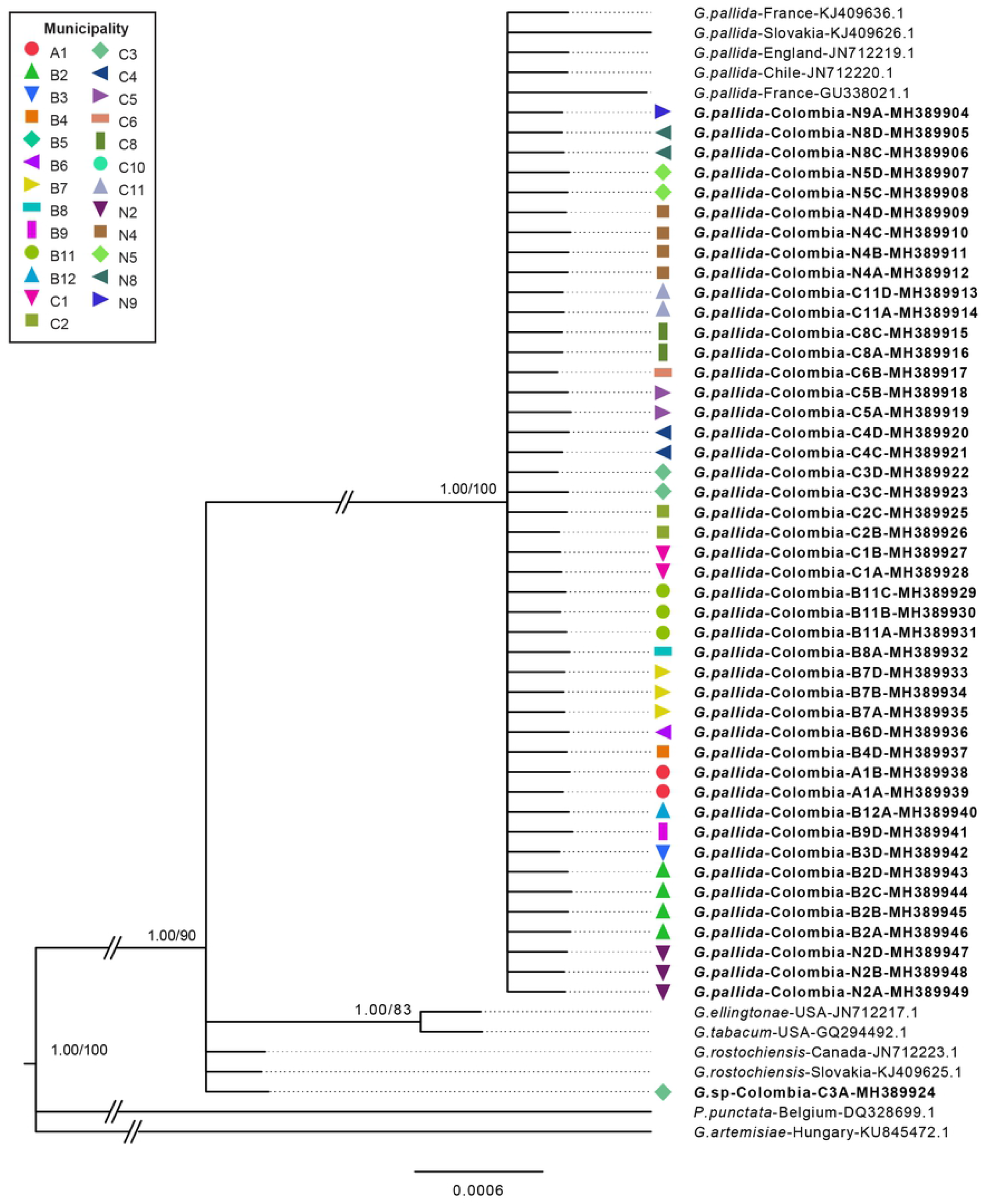
Phylogenetic relationships within the genus *Globodera*. Bayesian 50 % majority rule consensus trees as inferred from D2–D3 expansion segments of 28S rRNA as a single partition with HKY model. Node-support values: Left value posterior probability BI shown if >95%, right value bootstrap from ML analysis shown only if >70%. Newly obtained sequences in this study are in bold.

The 50% majority-rule BI consensus tree of the alignment generated for the 69 sequences of the region conformed by the internal transcribed spacer 1 and 2 including the 5.8S rRNA regions, showed two well supported major clades (PP=1.00/ BP=100) that were consistent with the findings based on 28S rRNA phylogeny (Fig 3). Clade (i) was formed by *G. pallida* and *G. mexicana*, and Clade (ii) was formed by *G. rostochiensis*, *G. tabacum*, *G. ellingtonae* and one sequence from Colombia. In Clade (i) two sequences of *G. pallida* from Peru along with the rest of sequences from Colombia formed a well-supported subclade (PP=1.00, BP=90) that was clearly separated (PP=1.00, BP=94) from one sequence of *G. pallida* from Peru. The sister clade of this sub-clade was formed by other sequences of *G. pallida* from Peru and European countries such as Ukraine, England and Poland, that were separated with high support (PP= 1.00, BP= 96) from sequences of *G. mexicana* and moderately support (PP= 1.00, BP= 79) of *G.pallida* from Chile and Peru. In Clade (ii) a single sequence from Colombia formed a well-supported subclade (PP=1.00, BP=93) along with *G. rostochiensis* from USA, Australia, Canada, England and Bolivia that was related with *G. tabacum* from Japan, USA and Argentina. This subclade formed a sister clade with *G. ellingtonae* from USA and Chile with high support (PP=1.00, BP=74).

**Fig 3.**
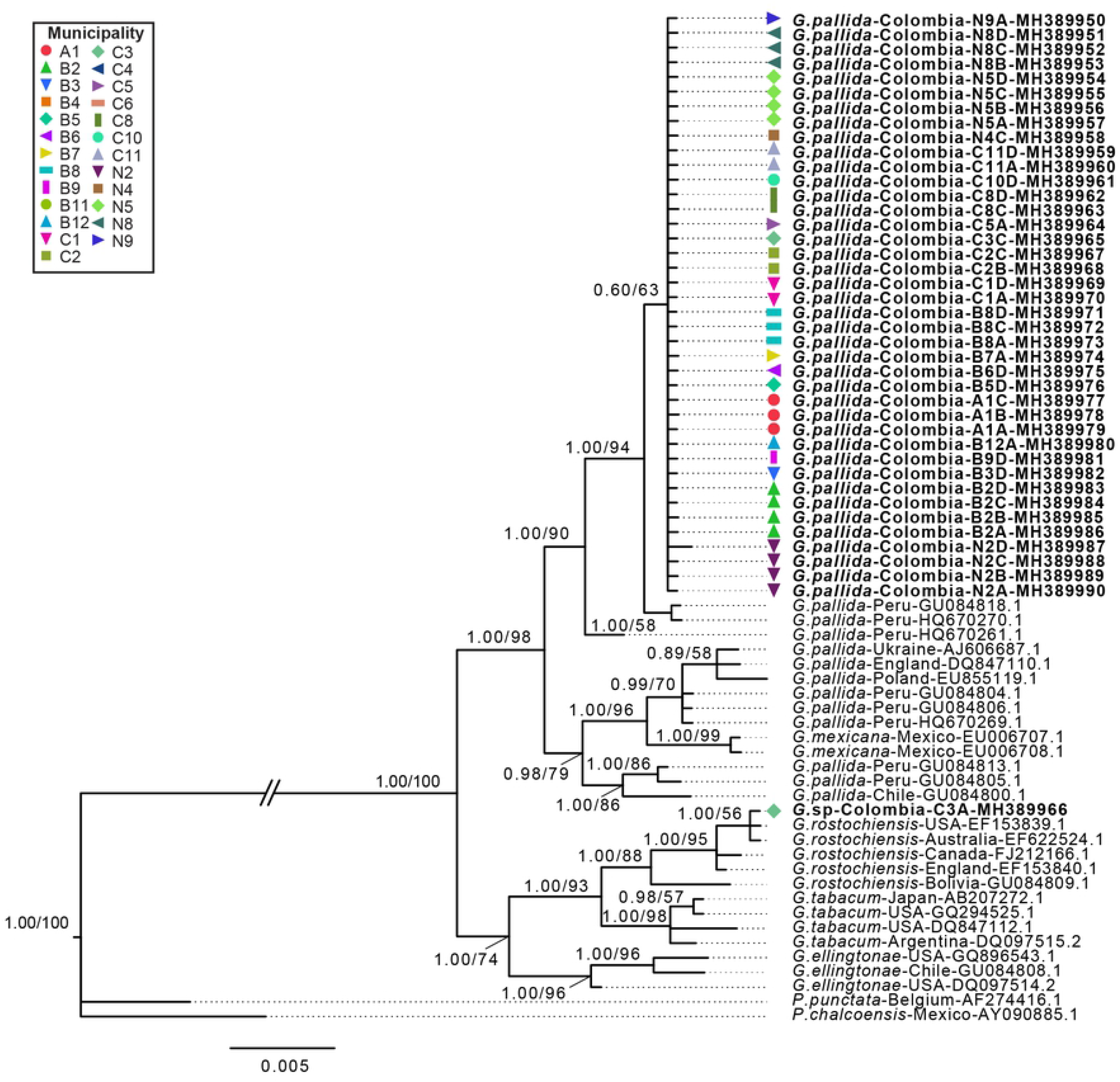
Phylogenetic relationships within the genus *Globodera*. Bayesian 50 % majority rule consensus trees as inferred from Internal Transcribed Spacer 1 and 2 plus 5.8S rRNA gene with first and second position with K80 substitution model, and a second partition with the third codon under K80+I model. Node-support values: Left value posterior probability BI shown if >95%, right value bootstrap from ML analysis shown only if >50%. Newly obtained sequences in this study are in bold.

Genetic distances among cyst nematodes sequences from Colombia and other nematodes species included in the phylogenetic analyses are summarized in Table 4. Based on the 28S rRNA gene sequences, all sequences from Colombia from Clade (i) had the lowest average number of nucleotide substitutions per site (Nucleotide divergence - *Dxy*= 0.002) and lowest number of net nucleotide substitutions per site (net genetic distance - *Da*= 0.001) when compared to *G. pallida* sequences from the other countries without fixed differences among groups (Table 4). The cyst nematode sequence from Colombia in Clade (ii) had the lowest divergence when compared with *G. rostochiensis (Dxy*= 0.001, *Da*= 0.000) without showing any fixed differences among groups (Table 4). In agreement with the 28S rRNA marker, the genetic distances based on the internal transcribed spacer 1 and 2 including the 5.8S rRNA gene sequences of cyst nematodes from Colombia in Clade (i) was lowest when compared with *G. pallida (Dxy*= 0.014, *Da*= 0.008), with one fixed substitution (Fig 3, Table 4) and lowest in Clade (ii) when compared with *G. rostochiensis (Dxy*= 0.003, *Da*= 0.000) with no fixed differences among groups (Table 4).

**Table 4.**
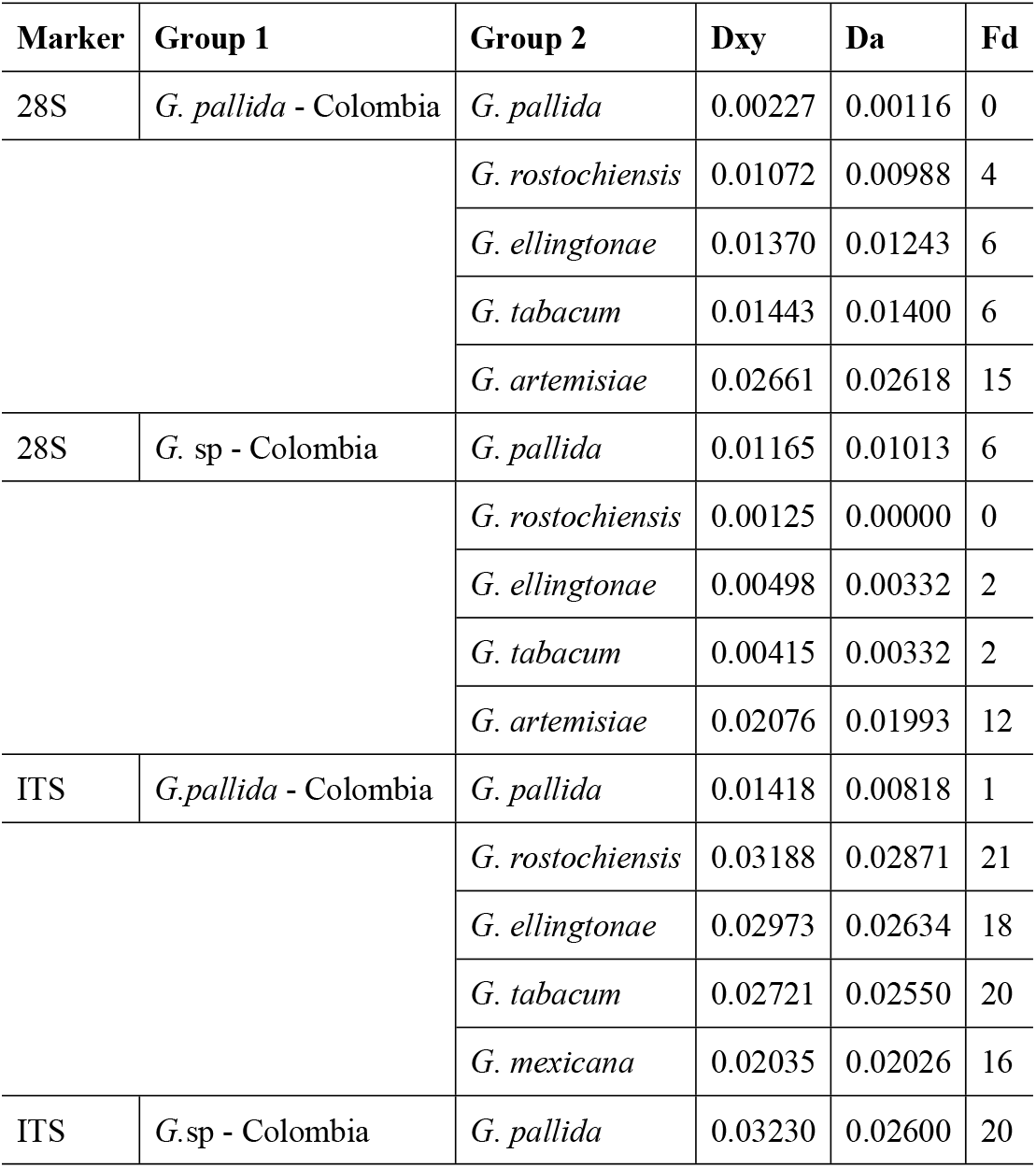
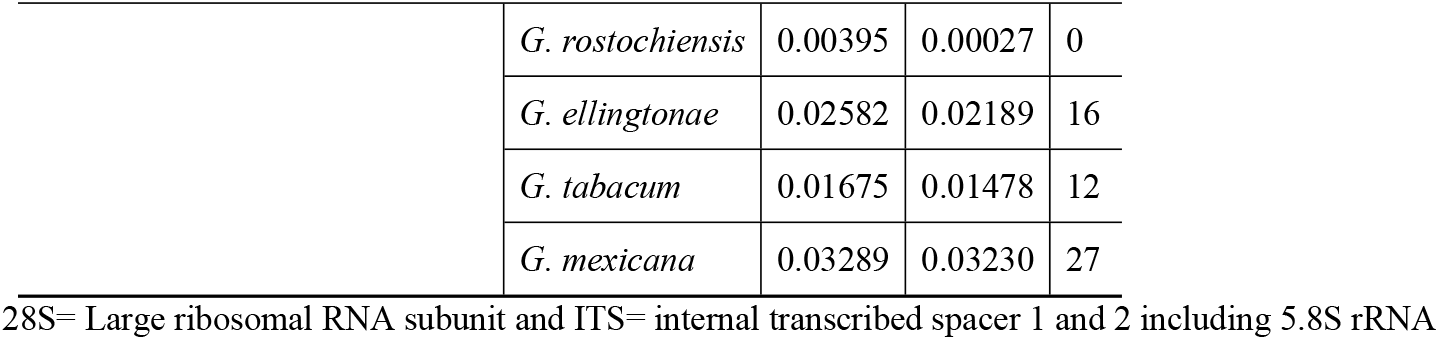
Gene divergence between potato cyst nematodes from Colombia and other species retrieved from GenBank. Dxy, Da and Fd correspond to average number of nucleotide substitutions per site, the number of net nucleotide substitutions per site, and number of fixed differences between compared groups, respectively.

### Cyst nematodes morphometric characterization

A total of 233 cysts were examined for morphometric measurements. Considerable degree of overlap was observed and all morphometric characters examined were significantly variable among populations (Fig 4): BL (DF= 24, F= 7.24, *P* < 0.0001), BW (DF= 24, F= 7.23, *P*= 0.0001), L/W ratio (DF= 24, F= 2.21, *P*= 0.0016), DAF (DF= 24, F= 2.39, *P*= 0.0005), FL (DF= 24, F= 3.23, *P* < 0.0001), GR (DF= 24, F= 3.23, *P*= 0.0001) and NCR (DF= 24, F= 1.68, *P*= 0.0297) (Fig. 4). Cysts in C10, showed the highest BL (639.42 ± 20.99 μm) and BW (637.05 ± 21.42 μm) and A1 population had the smallest mean size (417.67 ± 20.99 μm BL, 402.34 ± 21.43 μm BW). C3 showed significant differences on morphometric characters in the perineal area in comparison with the other populations, with a highest mean DAF (62.86 ± 5.43 μm), a lowest significant FL (18.43 ± 1.91 um), a significant higher GR (3.6) and the highest NCR (ranging from 10 to 16). For the other populations, DAF was in average 47.48 μm, FL was 26.45 μm, GR (1.8) and the NCR was 9. Nevertheless, some populations showed values outside of the range, for instance, B7 and B8 had the lowest mean DAF (36.28 ± 5.15 μm and 36.75 ± 5.43 μm) and B2 had a highest significant FL (36.30 ± 1.81 μm) and lowest GR (1.3) (Fig 4).

**Fig 4.**
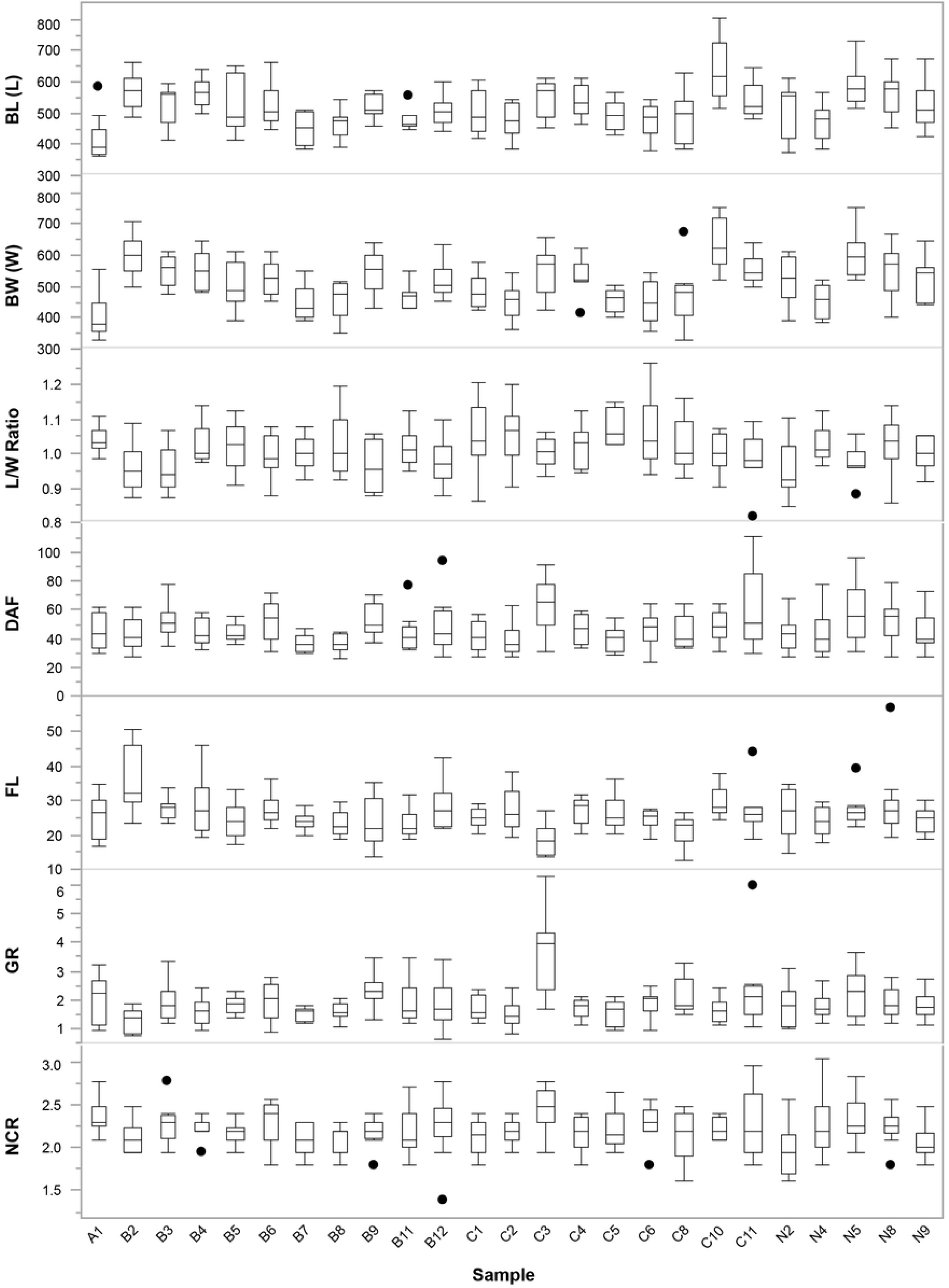
Boxplots showing morphological characters variation among PCN populations from cultivated potatoes in Colombia. Mean values shown for each population based on ANOVA (Tukey HSD test) at *P* ≤ 0.05). BL (Body length (μm): DF= 24, F= 7.24, *P* < 0.0001), BW (Body width (μm): DF= 24, F= 7.23, *P*= 0.0001), L/W ratio: DF= 24, F= 2.21, *P*= 0.0016, DAF (distance from anus to the nearest edge from fenestra (μm): DF= 24, F= 2.39, *P*= 0.0005), FL (Fenestra length (μm): DF= 24, F= 3.23, *P* < 0.0001), GR (Granek’s ratio: DF= 24, F= 3.23, *P*= 0.0001), NCR (Number of cuticular ridges: DF= 24, F= 1.68, *P*= 0.0297).

Two analyses were used to identify the variables that would separate species in the sampled populations. The LDA showed the variance associated with the first three canonical variables was 79% of the total variation and had the highest partial correlations between the canonical variables and the covariates, adjusted for the group variable (Table 5). Thus, the variables BW, NCR and GR in the SDA were selected in this order (Table 6). Canonical variable 1 had the highest correlation with BW (0.9392) followed by GR (0.2305), suggesting that separation of the groups on this axis was mostly due to plastic differences in body size. Canonical variable 2 had the highest correlation with GR (0.7752) followed by NCR (0.1785), therefore, separation of groups on this axis was mainly to differences in characters associated with features in the perineal area. Canonical variable 3 was most correlated with NCR (0.9838) followed by GR (0.5881), characters often used to discriminate PCN species (Table 6). The grouping and separation of populations using these three canonical variables is shown in Figure 5. Despite significant overlap of the 95% confidence ellipse for each population mean, canonical variable 1 distinctly separated some populations pairs with populations A1 and C10 at opposite sides of the axis. Canonical variable 2 distinctively separated population C3, which 95% confidence ellipse only partially overlapped with population C5 (Fig 5). Interestingly, few individuals from populations C11, B9, N2 and, C8 were also near population C3 95% confidence level ellipse mean, as few C3 individuals were more similar to other populations 95% confidence level ellipse mean (Fig 5).

**Table 5.**
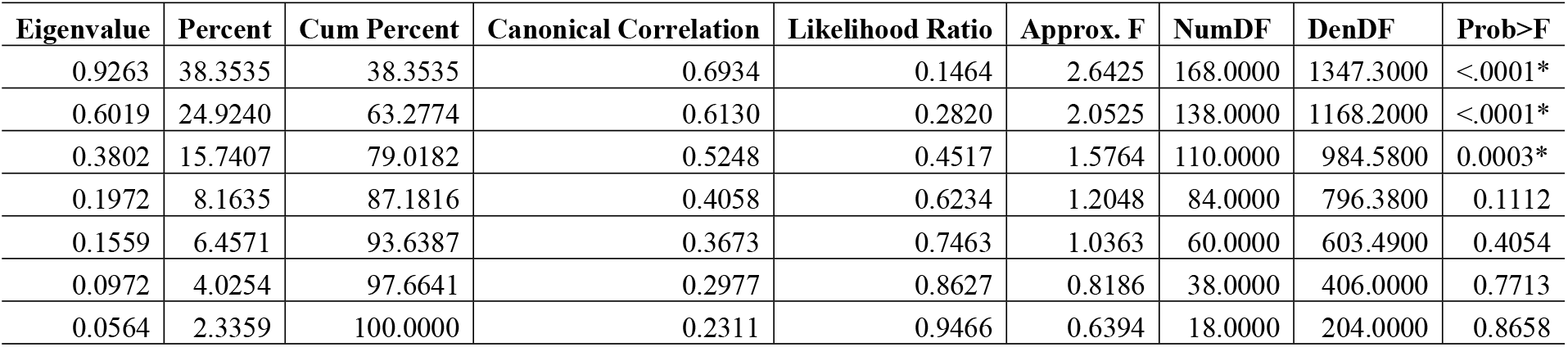
Canonical details calculated from the overall pooled within-group covariance matrix of potato cyst nematodes morphometric characterization.

**Table 6.** Canonical structure and standardized canonical coefficients of potato cyst nematodes morphometric characterization.

| Variable* | Pooled within canonical structure |  |  | Pooled within-class standardized canonical coefficients |  |  |
| --- | --- | --- | --- | --- | --- | --- |
|  | Can1 | Can2 | Can3 | Can1 | Can2 | Can3 |
| BW | 0.9392 | -0.2590 | 0.2255 | 0.9663 | -0.3197 | 0.0428 |
| NCR | 0.0155 | 0.1785 | 0.9838 | -0.4807 | -0.7352 | 1.1574 |
| GR | 0.2305 | 0.7752 | 0.5881 | 0.4335 | 1.3524 | -0.2521 |

\* BW= body width; NCR = Number of cuticular ridges between vulva and anus; GR = Granek's ratio.

**Fig 5.**
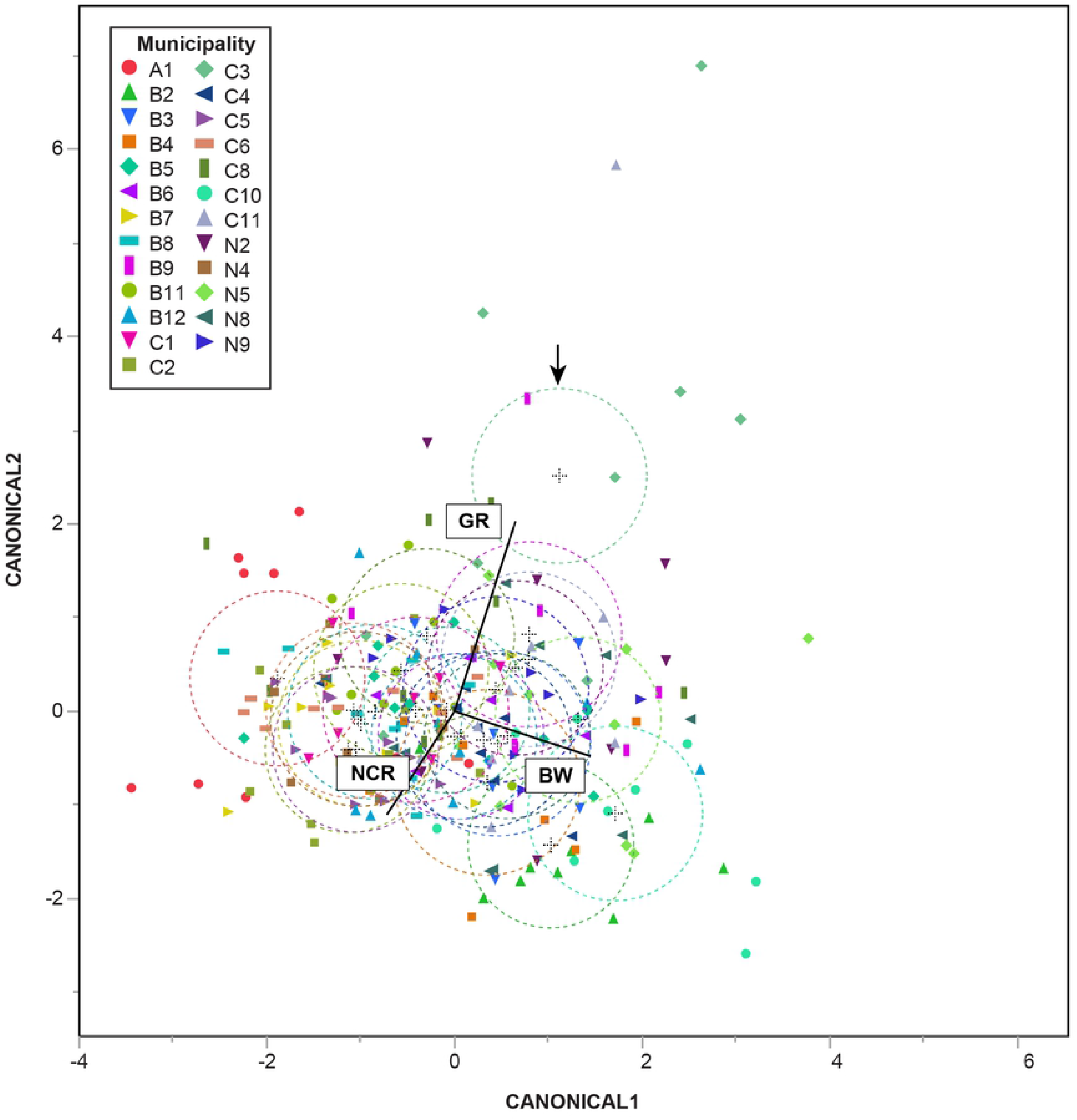
Canonical plot scores and 95% confidence ellipses from stepwise discriminant function analysis of three morphometric characteristics of PCN nematode cyst from Colombia. Each ellipse corresponds to a 95% confidence limit for the population multivariate mean. Significantly different populations have non-intersecting circles. The set of rays that appears in the plot represents the covariates in which the length and direction of each ray indicates the degree of association of the corresponding covariate with the first two canonical variables. The arrow indicates C3 population 95% confidence interval. NCR, number of cuticular ridges, GR, Granek’s ratio, BW, body weight.

## Discussion

Even when PCN is considered a re-emerging potato pathogen in Colombia, first identified in the department of Nariño in 1970 [23], documented surveys only report PCN in municipalities of Nariño, Cauca, Boyacá and Cundinamarca [24,25]. Our comprehensive study that sampled 75 municipalities in 9 potato producing departments of Colombia found that 60% of the tested samples were positive for PCN (355 out of 589 sampled fields), and that the pathogen is widespread in all Colombian producing potato departments (Cundinamarca, Boyacá, Nariño, Antioquia, Cauca, Norte de Santander, Santander, Tolima and Caldas), with cysts that contain viable eggs present in all sampled departments.

However, there was variation in population densities among departments. The highest densities were found in Cundinamarca and Boyacá, ranging from 1 to 1,327 cyts/100 g of soil and 1 to 873 cysts/100 g of soil, respectively. Nieto et al. (1983) [24] surveyed these departments in early 1980s, but since detected cysts were empty or contained non-viable eggs, these regions were declared as PCN free. Later, Arciniegas et al (2012) [25], reported PCN in Tunja, Samacá and Ventaquemada in Boyacá and Tausa, Tabio and Zipaquirá in Cundinamarca. In our study, PCN was detected in new municipalities with the highest densities found in Tausa, Ubaté and Villapinzón in Cundinamarca and, in Arcabuco, Belén, Tunja, Samacá, Sogamoso and Sora in Boyacá (Table 1). Boyacá and Cundinamarca are the largest producers of potatoes in Colombia, with 40,724 and 61,322 harvested hectares corresponding to 26% and 39% of the total potato national production in 2018, respectively [26]. Potatoes are the main plant crop grown by farmers and fields are usually planted in monocultures for several continuous cycles. As PCNs are highly specialized, sedentary and obligate endoparasites of solanaceous plants [1,22,54], the constant presence of potato crops in monocultures for several cycles may lead to persistence and increase of this plant pathogen in these two regions overtime. In contrast, in Nariño department, although the nematode was found in all municipalities sampled, PCN densities were lower, from 1 to 345 cysts/100 g of soil, and similar ranges were found by Nieto et al (1983) [24]. Nariño ranks third in production with 31,611 harvested hectares (19,35% of total potato national production), in contrast to Boyacá and Cundinamarca, in this department farmers usually grow different potato cultivars within a field (e.g. Diacol Capiro, Pastusa Suprema, Betina) with a crop rotation scheme usually with non-host plants such as corn, cabbage, lettuce, onion and pastures, and the use of biological microorganisms for the control of other pest problems has also implemented [55], which reduces pesticides use. Similar potato production scheme was observed in its neighbor department, Cauca, and PCN densities decreased in the latter from 23 cysts/100 g of soil [24] to 5,97 cysts/100 of soil in this study. Considering that *G. pallida* requires a living potato plant to complete its life cycle [6], the management practices implemented in both departments may reflect the reduction of PCN densities in these regions.

Our results also show *G. pallida* populations have spread into new regions of Colombia. In the departments of Antioquia, Caldas, Tolima, Santander and Norte de Santander, PCN was detected in all municipalities sampled, although with low population levels (5.97 cysts/100 of soil in average in Antioquia, 0.95 cysts/100 of soil in Norte de Santander, 0.8 cysts/100 of soil in Santander, 0.43 cysts/100 of soil in Caldas and 0.3 cysts/100 of soil in Tolima). To our knowledge, this is the first report of the presence of *G. pallida* in these departments. These regions represent in general, low potato growing areas with low participation in national potato production (2.27% in average in 2018) [26], and were therefore considered before as PCN free. The spread of PCN is mainly caused through tubers, soil or equipment contaminated with cysts [54] and potato seed tubers in these departments frequently come from Boyacá and Cundinamarca, which may allow the dissemination of this plant pathogen into these new regions. Nevertheless, population levels in these departments are low and the extent of PCN is limited, therefore, to maintain low levels and to avoid the spread into new areas, intensive monitoring program for PCN should be implemented in all potato producing regions of Colombia.

### Molecular identification and phylogenetic relationships of PCN species from Colombia

The molecular phylogeny of PCN populations based on ITS1-5.8S-ITS2 rDNA and 28S D2-D3 regions supported the presence of at least two PCN species in Colombia. *Globodera pallida* was found in all populations that resulted positive for PCN and, molecular phylogeny based on the ITS1-5.8S-ITS2 rDNA grouped all *G. pallida* from Colombia in a single clade that was closely related to P5A pathotype strains (GenBank accession number HQ670270.1) and clone La Libertad (GenBank accession number GU084818) (Fig 3). This finding suggests that *G. pallida* present in Colombia have a different origin than *G. pallida* present in countries such as Ukraine, England and Poland that cluster as a monophyletic clade with other Peruvian strains, and *G. pallida* present in Chile that cluster with a different Peruvian strain (Figs 2 and 3). Despite that P5A peruvian strain has been considered as a different species [12,17], a recent study based on ITS rRNA, *COI* and *cytb* mitochondrial regions concluded that all clades within *G. pallida* belong to a single species [2]. The 28S D2-D3 phylogeny, although with lower level of resolution, also clustered all but one PCN populations from Colombia with *G. pallida* around the globe as a monophyletic clade (Fig 2). For both gene regions, a single sequence from the C3 population grouped in a distant clade along with individuals of *G. rostochiensis*. Genetic distance analyses based on gene regions ITS1-5.8S-ITS2 and D2-D3, were congruent with these findings showing *G. pallida* from Colombia with the smallest *Da* (0.8 % and 0.12% for ITS rDNA and D2-D3, respectively) and the smallest *Dxy* (1.41 % and 0.23 % for ITS rDNA and D2-D3, respectively) when compared with other *G. pallida* populations. Similarly, genetic distances from C3 population showed the lowest distance when compared with *G. rostochiensis* (*Dxy*= 0.001, *Da*= 0.000).

Therefore, the ITS1-5.8S-ITS2 rDNA and 28S D2-D3 molecular analyses were able to identify with high phylogenetic support *G. pallida* and *G. rostochiensis*. Additionally, ITS1-5.8S-ITS2 rDNA phylogenetic resolution supports a northern Peru origin of *G. Pallida* present in Colombia, nevertheless this hypothesis must be further investigated using additional samples and molecular markers, with statistical inference such as model testing and coalescent demographic reconstruction. Although with less taxa included, molecular phylogeny based on 28S D2-D3 gene improved the node support found in previous phylogenies between *G. pallida* and *G.tabacum* (i.e. PP = 54 and 72%) (e.g.,[9,16]), and the unresolved positions for *G. rostochiensis* [16]. Taken all together, both DNA markers used in this study showed to be useful to identify *Globodera* species present in Colombia, with ITS1-5.8S-ITS2 rDNA being more informative in phylogeographic perspective [12].

### Morphometric identification of PCN species from Colombia

The use of stepwise and canonical analysis is an effective method for grouping and distinguishing species and populations from different taxa [56–59]. Despite a significant degree of overlap in morphometric data (Figs 4 and 5), canonical discriminant analyses identified two main groups based on cyst morphometric measurements. *Globodera pallida* individuals were observed in the canonical discriminant plot as populations were overlapping across canonical 1, indicating a wide variation in BW size associated with phenotypic plasticity (Fig 5). The wide morphometric variation observed within *G. pallida* from Colombia populations have also been reported in other populations of *G.pallida* as well as for other *Globodera* species [9,17,60]. It is often expected that observed morphological variation within *Globodera* species increases as new populations are analyzed [7,17]. However, since wide morphometric variability within and among populations of *G. pallida* was detected, future research should be performed to elucidate the degree in which cryptic species and phenotypic plasticity occur among PCN associated with potato crops in Colombia. On the other hand, individuals from C3 population were observed in the top portion of the canonical discriminant analyses, with all but one individual outside the C3 95% confidence ellipse across canonical 2 (Fig 5). The presence of individuals from population B9, N2 and C11 near population C3 position in the canonical space, suggests the possibility of a wider distribution of *G. rostochiensis* associated Colombian potato crops, since morphological measurements of these individuals fit in the range of *G. rostochiensis*. However, to confirm the presence of this PCN species in Colombia, further research should include genetic and morphological data from additional individuals (cysts, juveniles and males) as well as pathogenicity assessment of these Colombian isolates.

## Conclusions

This study provides new information about the status and prevalence of PCN species associated with cultivated potatoes in the main producing regions of Colombia including for the first time genetic and morphological information. Molecular phylogenies with ITS1-5.8S-ITS2 rDNA and D2/D3 28S regions and morphometric measurements of cysts were effective in the identification of *G. pallida*, the dominant species present in all departments surveyed in this study, and suggest the presence of *G. rostochiensis*, in one municipality of Cundinamarca, which is currently under description. Considering the presence of PCN species constitute a threat for potato production, intensive sampling and monitoring of this plant pathogen should be conducted in order to reduce and prevent the spread into new areas. The development of management practices that involves the evaluation of resistant varieties for populations that tested positive for PCN, as well as other practices such as crop rotations, trap crops, biofumigants, biocontrol agents among others that have shown to be effective for other *G. pallida* populations worldwide, are also a crucial step to reduce population densities of PCN in Colombia.

## Acknowledgments

The authors thank the Colombian Ministry of Agriculture for funding the Project “Recomendaciones técnicas para el manejo integrado de los problemas fitosanitarios: *Globodera pallida*, síndrome X, virus PYVV y sus posibles vectores en papa”, as well as the supporting research assistant of Corporación Colombiana de Investigación Agropecuaria, La Selva Research Station, Mario Alonso Mesa, for greenhouse and field work.

## Contributions

**Conceptualization:** Olga Y. Pérez, Claudia M. Holguin.

**Data curation:** Daniela Vallejo, Diego A. Rojas, John A. Martinez, Sergio Marchant, Claudia M. Holguin, Olga Y. Pérez.

**Formal analysis:** Sergio Marchant, Claudia M. Holguin, Daniela Vallejo.

**Investigation:** Daniela Vallejo, Diego A. Rojas, John A. Martinez, Sergio Marchant, Claudia M. Holguin, Olga Y. Pérez.

**Methodology:** Daniela Vallejo, Diego A. Rojas, John A. Martinez, Sergio Marchant, Claudia M. Holguin, Olga Y. Pérez.

**Writing - original draft:** Claudia M. Holguin, Sergio Marchant, Daniela Vallejo, Diego A. Rojas, John A. Martinez, Olga Y. Pérez.

**Writing - review & editing:** Claudia M. Holguin, Sergio Marchant, Olga Y. Pérez.

